# Minimum Error Calibration and Normalization for Genomic Copy Number Analysis

**DOI:** 10.1101/720854

**Authors:** Bo Gao, Michael Baudis

## Abstract

Copy number variations (CNV) are regional deviations from the normal autosomal bi-allelic DNA content. While germline CNVs are a major contributor to genomic syndromes and inherited diseases, the majority of cancers accumulate extensive “somatic” CNV (sCNV or CNA) during the process of oncogenetic transformation and progression. While specific sCNV have closely been associated with tumorigenesis, intriguingly many neoplasias exhibit recurrent sCNV patterns beyond the involvement of a few cancer driver genes.

Currently, CNV profiles of tumor samples are generated using genomic micro-arrays or high-throughput DNA sequencing. Regardless of the underlying technology, genomic copy number data is derived from the relative assessment and integration of multiple signals, with the data generation process being prone to contamination from several sources. Estimated copy number values have no absolute and linear correlation to their corresponding DNA levels, and the extent of deviation differs between sample profiles which poses a great challenge for data integration and comparison in large scale genome analysis.

In this study, we present a novel method named Minimum Error Calibration and Normalization of Copy Numbers Analysis (*Mecan4CNA*). For each sCNV profile, *Mecan4CNA* reduces the noise level, calculates values representing the normal DNA copies (baseline) and the change of one copy (level distance), and finally normalizes all values. Experiments of *Mecan4CNA* on simulated data showed an overall accuracy of 93% and 91% in determining the baseline and level distance, respectively. Comparison of baseline and level distance estimation with existing methods and karyotyping data on the NCI-60 tumor cell line produced coherent results. To estimate the method’s impact on downstream analyses we performed GISTIC analyses on original and *Mecan4CNA* data from the Cancer Genome Atlas (TCGA) where the normalized data showed prominent improvements of both sensitivity and specificity in detecting focal regions.

In general, *Mecan4CNA* provides an advanced method for CNA data normalization especially in research involving data of high volume and heterogeneous quality. but with its informative output and visualization can also facilitate analysis of individual CNA profiles. *Mecan4CNA* is freely available as a Python package and through Github.

## Introduction

Copy number aberrations (CNA^1^) represent the regional gain and loss of DNA copies of chromosome regions. Somatic copy number aberrations are frequently observed in cancer tumors and they exhibit complex and intrinsic connections to the development of cancer [3]. CNA data is widely used in both clinical practice and research. The evaluation of a patient’s CNA profile has become a common procedure in treatment for certain cancer types. By analyzing CNA datasets, researchers have been able to identify and verify mutational patterns and genes associated with CNAs in various cancer types [25]. With the rapid advance of technology and the explosion of data, more subtle driver genes and more comprehensive patterns are expected to be revealed in the near future [10, 21]. Although CNA data has become a principal component in cancer genome analysis, the interpretation of CNA data is still a challenging task. This is primarily due to the noise contamination that is prevalent among CNA data.

Currently, CNA data is usually generated using either one of several micro-array technologies [19, 15, 26] or increasingly utilising high-throughput sequencing methodologies [22, 24]. Independent of technology or specific platform, CNA data often has a high noise level that can be attributed to three main sources. First, the noise level is heavily influenced by the purity of the tumor sample. Ideally, a copy number should always be an integer value and correspond to a DNA level of the measured tumor cells. In practice, the tumor sample is often a mixture of normal cells, main tumor cells, and minor tumor clones with partially diverging mutation profiles, resulting in overall and possibly regional divergences from the copy number levels expected from a homogeneous tumor sample with the measured DNA levels potentially representing averages of different CN states. Second, the CN value is derived from the relative measurement of DNA content, with non-linear correlation between the true regional CN and the measured – intensity or count based – signal. Third, systematic errors accumulate in each experiment step throughout the pipeline, e.g. in sample preparation or labelling procedures. In theory, the sources of experimental noise can be alleviated by performing additional experiments, increasing source material use or sequencing depth or fine tuning experimental and analytical protocols. However, such efforts are counteracted by scarce or inherently limited (e.g. archival tissue) source material, high cost (e.g. technical replicates or increased read depth) or analytical overhead. Most importantly, experimental improvements cannot be pursued in meta-analyses which rely on pre-existing data, usually derived from a variety of sources.

The frequently high noise level in CNA data leads to two common problems in downstream analysis procedures. First, the value of the copy number level in a genomic region usually reflects a log2 normalised relative deviation instead of the actual integer DNA level of the main tumor cell clone. Second, the extent of the deviation differs between profiles of samples with the same underlying CNA value. To accommodate for these problems, CNA data needs to be calibrated (i.e. value adjusted) in single profile analysis, and to be normalized for the analysis of multiple profiles. Currently, there is no “Gold Standard” methodology, for neither the calibration nor the normalization of CNA data. The most common approach is manual assessment, it works well for a single profile with a simple aberration pattern, but becomes incompetent when facing a complicated pattern or a dataset. Another widely used approach is median centering, it is intuitive and easy to implement, but it has been shown that this method also struggles when dealing with complex profiles [7, 6]. In clinical evaluation, contextual information and additional data are usually used to make a good assessment. But this approach depends on expert knowledge and extra data, it is not applicable for most research settings. Finally, several computational methods were developed in the last decade to detect tumor purity or actual DNA levels [1, 2, 5, 6, 7, 8, 13, 14, 23]. Typically, ABSOLUTE [6] uses allele specific copy number ratio and pre-computed models to estimate purity and ploidy, and from that computes potential models of absolute copy numbers. BACOM [7] exploits allele specific copy number signals with a Bayesian model to differentiate homozygous and heterozygous deletions and estimate normal cell fraction. AbsCN-seq [13] uses a statistical method to infer purity, ploidy and absolute copy number. These methods can provide more reliable calibrations in complex scenarios. Unlike in calibration, where methods are usually generic, normalized methods are often study specific [4, 9, 16]. In some copy number studies, where a large number of CNA data were compared for analysis, the data was even not normalized but merely calibrated. To our knowledge, there exist no generic normalization methods for CNA data at the moment. Although the computational methods for calibration can be extended to perform normalization, they bear a few drawbacks. First, most of these methods are platform specific, which limits the scope of their application. Second, they often require the original raw file, or a control sample, or both. These files are not always available. Third, they do not provide direct functions for normalization. Therefore, additional works are needed, which often involves complex scripting and sometimes manual tuning. Last, these methods usually are based on complicated models that invoke heavy computational demand. It makes them both time and resource consuming when processing high volume datasets.

In this study, we present Minimum Error Calibration and Normalization for Copy Numbers Analysis (Mecan4CNA), a generic computational method aimed to address the aforementioned issues. Instead of estimating the exactly purity or DNA levels, *Mecan4CNA* transforms the problem to the estimation of the baseline (the level of normal DNA copies) and the value distance between DNA levels. Then *Mecan4CNA* computes these two values for each profile and uses them to normalize data uniformly. All CNA profiles will be normalized to the same scale, where copy number values actually correspond to their true DNA levels. *Mecan4CNA* can be used on CNA data generated from any platform and pipeline. It only requires segmentation files, which is the final product of all copy number pipelines. *Mecan4CNA* is based on an algebraic model, which makes it efficient both in speed and resource. Although *Mecan4CNA* is primarily designed for data normalization, it can also be used for quick calibration. It provides detailed information and visualization on estimation results, which can be used for a more in-depth analysis of each profile. *Mecan4CNA* is freely available as a Python package and through Github.

## Methods

A copy number value in a CNA profile represents the number of DNA copies (alleles) in the corresponding genomic region. As a general description, a CNA profile and the value of a specific position can be modelled as the following:

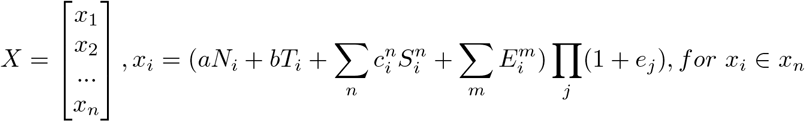

where X represents a CNA profile of n positions, and *x_i_* represents the value at a specific position. *N_i_*, *T_i_* and *S_i_* denote the actual DNA copy numbers of normal cells, tumor cells and sub-clone cells, respectively. Likewise, *a, b* and *c* are the composition ratios of normal cells, tumor cells and sub-clone cells at this position, respectively. 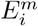 represents an independent error of this position and *e_j_* represents a systematic error for the entire sample. For each *x_i_*, 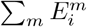 and Π_*i*_(1 + *e_j_*) are constants. If we also consider the summation of sub-clones as a single pseudo-sub-clone, *x_i_* can be simplified as the following:

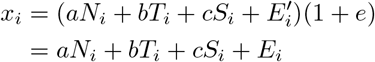

where *a, b, c* and *e* are constants in each sample. *N_i_, T_i_, S_i_* and *E_i_* are variables of each *x_i_*. Here, *E_i_* represents the integral of all errors.

Ideally, the composition and noise level of a sample can be retrieved by solving matrix *X*. In practice, all parameters except *x_i_* are unknown, thus making the solution of *X* a daunting task. In previous studies, statistical models and information of controls were used to circumvent this problem. Here, we presents an algebraic approach that requires no additional information. The fundamental principle is to cancel out parameters through operations and aim for a partial solution. Let *D*(*i, j*) be the distance of *x_i_* and *x_j_*:

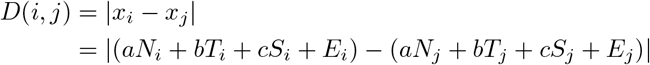

Because germline copy number changes are rare events and are usually filtered by most processing pipelines, we can assume *N_i_* to be a constant of 2. Therefore, *D*(*i, j*) can be simplified as:

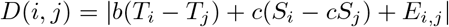

Let *R*(*i, j, k*) represents the ratio of two distance *D*(*i, j*) and *D*(*i, k*)

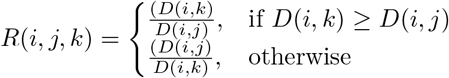

Here, we illustrate the deduction of one case, and the other scenario is the same.

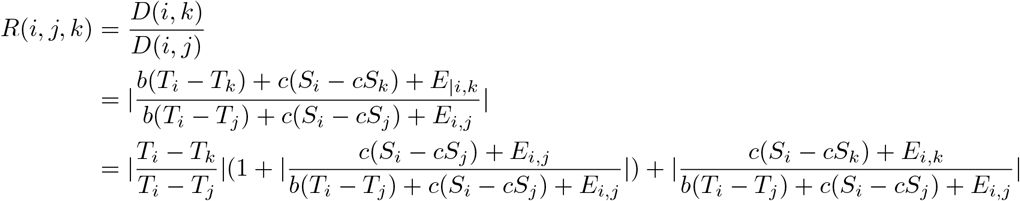

Here, 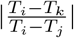 is the ratio of the distances between actual copy number levels. When *b* >> *c*, which means a biopsy where the proportion of the main tumor is much higher than the proportion of sub-clones, we can have:

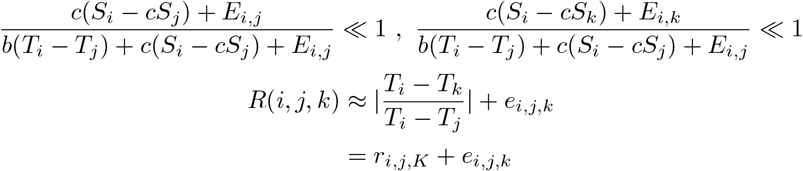

Because *e_i,j,k_* reflects the deviation from the actual copy number, it is smallest when *x_i_, x_j_* and *x_k_* are all the corresponding values of actual DNA levels. If we can find a pair *x_i_* and *x_j_* that has the lowest overall deviation, we can say that *x_i_* and *x_j_* represent two DNA levels in a solution of *X*. And that is to calculate the following equation:

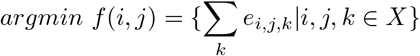

Furthermore, if *T_i_* and *T_j_* are two adjacent DNA levels, then |*T_i_* − *T_j_*| = 1 and *r_i,j,k_* becomes an integer. In most cases, when the data is processed properly, the systematic error term *e_i,j,k_* is relatively small. Therefore, we can evaluate *e_i,j,k_* as in the following:

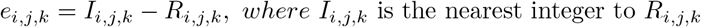

Interestingly, the systematic errors of a CNA profile are not uniformly distributed. Usually, positions that are close to the normal level have lower errors and positions far away from the normal level tend to have higher errors. Therefore, the *x_i_* and *x_j_* pair that has the lowest overall error sum are usually the normal level and one of its adjacent level.

Now, we have acquired a partial solution: a pair of *x_i_* and *x_j_* from *D*(*i, j*) which represents the distance between the normal DNA level and an adjacent level. We do not know which value actually corresponds to the normal level, and which corresponds to the adjacent level. By using a weighting function, we consider the strength of signal, the distance to the center and the validness of models comprehensively. One with a higher *B_score_* is determined as the baseline:

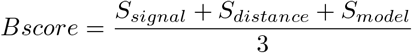

Theoretically, we have to evaluate *f*(*i, j*) for all pairwise combinations in *X*, which is a huge amount of computation. In practice, copy number signals are not randomly distributed. Most signals are normally distributed with a mean that corresponds to an actual copy number value. We only need to compute *f*(*i, j*) for these mean values, as they carry the lowest noise and are closest to the actual value. Furthermore, to reduce the complication of noise and small sub-clones, we choose a peak value only if it is covered by a minimum of p probes. For quick estimation, we assume 1% of the data is noise and *x_i_* distribute uniformly in an interval. When using a resolution of *n* bins in this interval, the probability density of each bin is 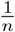. When there is a total of *m* noisy probes, 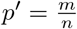, which is the expectation of a binomial distribution 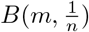. For an Affymetrix SNP6 array with approximately 2 * 10^6^ probes, if we take a resolution of *n* = 100, then *p*′ = 200. We want to choose a *p* that is greater than *p*′, so it can filter out noisy data and improve calculation speed. Empirically, we used *p* = 2000 as the threshold in our experiments.

## Results

In order to validate the usefulness of *Mecan4CNA*, first we compared the estimation results of our method with results from karyotyping and ABSOLUTE on cell line data. Additionally we evaluated the method on a series of simulated data, explored the baseline deviation situation of copy number data from TCGA, and finally used *Mecan4CNA*-normalized TCGA data as input for a GISTIC [12] analysis, to evaluate a possible improvement compared to using the unprocessed original data.

### Performance on real data

The NCI-60 tumor cell lines were selected by the National Cancer Institute of the United States (NCI) as a reference panel for drug screening experiments. It comprises 60 human cancer cell lines from 9 different cancer types. Their molecular characteristics have been well studied in the past two decades. Particularly, 58 cell lines have been karyotyped using the spectral karyotyping protocol [17], and the copy number variations of all cell lines have been profiled using microarray experiments [18]. DNA copy number changes on chromosome 13 were explored in detail in the karyotyping study. Among all karyotyped cell lines, 30 have either no changes or only chromosome level changes on chromosome 13. They represent ideal examples to validate the performance of our method, because we can match the karyotyping data with the microarray data to identify the copy number values (from microarrays) that correspond to the actual normal and abnormal DNA levels (from karyotyping). If we compare these values with the estimation results of *Mecan4CNA* on chromosome 13, we can evaluate the performance of the method.

When comparing data from microarrays with the karyotyped standard, 6 cell lines showed contradicting results and were excluded from further analysis (5 karyotyped as one copy loss but microarray showed normal; 1 karyotyped as one copy loss but microarray showed copy gain). These differences may be explained by a possible clonal evolution between the reference analysis and the cell line batches from which the microarray data was prepared. The remaining 24 cell lines were used for further analysis. In order to compare the performance of *Mecan4CNA* with existing methods, we also used ABSOLUTE, a widely used method to estimate purity and ploidy from copy number data, to infer the corresponding copy number levels of each cell line. Figure 1 shows the comparison results of *Mecan4CNA* and ABSOLUTE, respectively. On both graphs, the estimated values were plotted against the actual value of a DNA level. *Mecan4CNA* achieved 0.987 for spearman correlation coefficient and 0.033 for root mean square error (RMSE). Both scores indicate strong and confident correlation between the estimation and actual values. ABSOLUTE achieved 0.988 for spearman correlation coefficient and 0.049 for root mean square error (RMSE). This high concordance confirmed the solid performance of both methods on estimating copy number levels.

**Figure 1:**
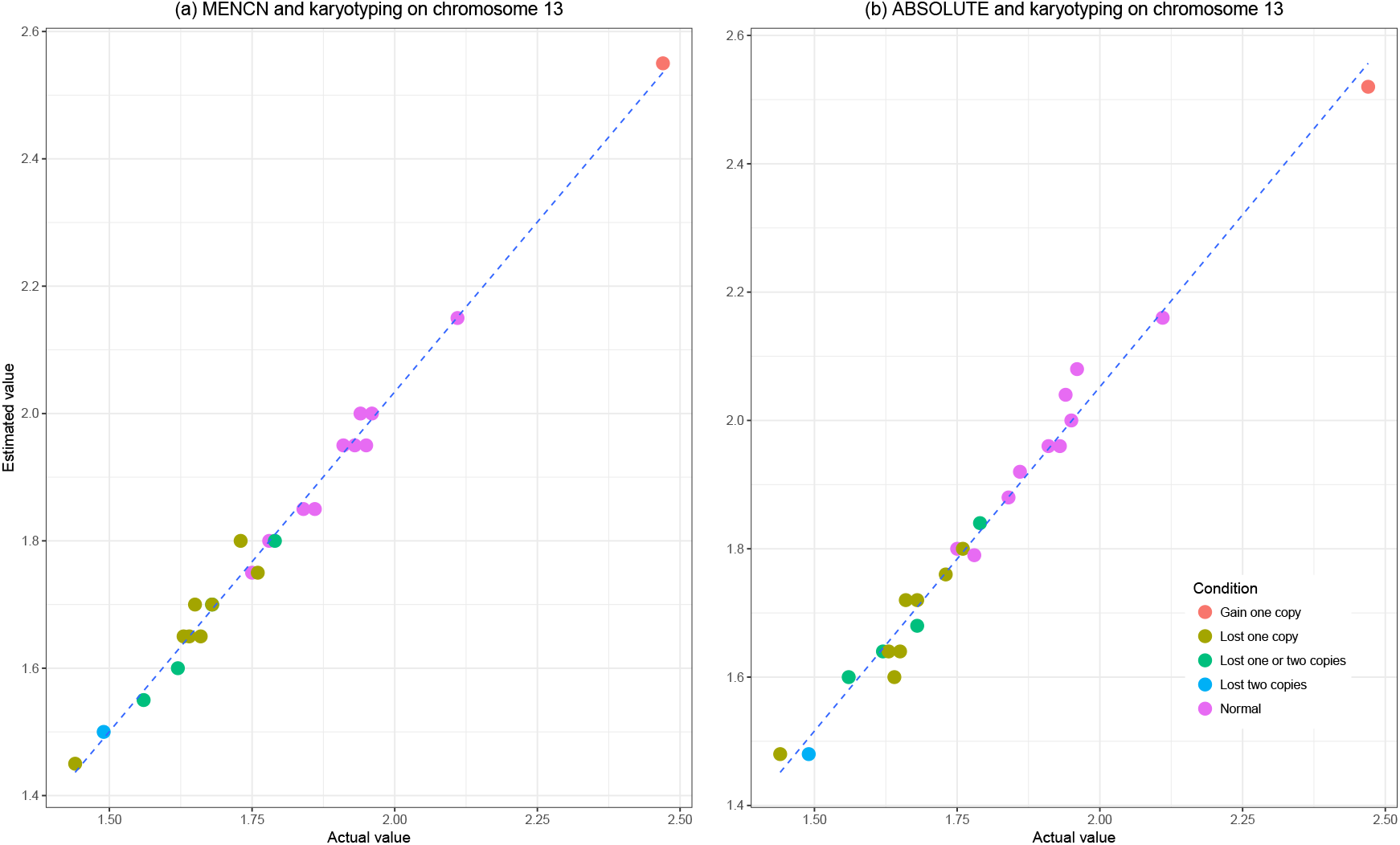
The comparison of estimated values and actual values on chromosome 13. The estimated values were calculated from the CNA file; the actual values were derived from karyotyping results. (a) estimation of *Mecan4CNA;* (b) estimation of ABSOLUTE. The estimation results of two methods showed high accuracy and coherence.

### Performance on simulated data

To further evaluate the performance of *Mecan4CNA*, we applied the method to 10 simulated datasets of different cell composition and noise levels. Every dataset comprises 100 samples, which are copy number segment profiles generated based on our modeling equation introduced in the method section. The cell composition was generated using a Dirichlet distribution. In high tumor composition samples, the composition ratio of normal and sub-clones are capped at 0.2. For both high tumor composition datasets and low tumor composition datasets, the independent errors were generated using a normal distribution with the standard deviation increasing from 0.001 to 0.009 with a step size of 0.002; the global errors were generated using a normal distribution with the standard deviation increasing from 0.1 to 0.18 with a step size of 0.02. The ploidy of normal cells was assumed to always be two. A total of 1000 samples were generated in this manner, and both the actual cell composition and tumor ploidy were known.

The five datasets containing samples with high contribution of the main (virtual) tumor clone show a gradual increase of noise levels from dataset to dataset. A similar observation can be made inthe other five datasets with low tumor composition (high sub-clone or normal).

As shown in Figure 2, the estimation accuracy of *Mecan4CNA* drops when the noise level increases or when the tumor composition decreases. Under the same noise level, for both the baseline and level distance, the estimation accuracy decreases significantly between high and low tumor composition. Under the same tumor composition, the estimation accuracy only shows slight declines when the noise level increases. The baseline estimation performed reasonably well even with low tumor composition and high noise level. However, the decrease of tumor composition has a greater impact on the estimation accuracy of level distance. Because even in low tumor composition, as long as the genome is not in complete chaos, which occasionally happens in cancer, the signal of normal DNA level (baseline) is often still detectable. But when the biopsy has a high composition of normal and sub-clone cells, the signal of the main tumor will be significantly reduced, and the signal of sub-clones becomes more influential at the same time. The combination of these two effects makes it much more difficult to have accurate estimation of main tumor’s level distance in samples with low tumor composition. *Mecan4CNA* provides conservative estimates of the level distance since overestimation can lead to undesired information loss during normalization, and underestimation only introduces false-positive information.

**Figure 2:**
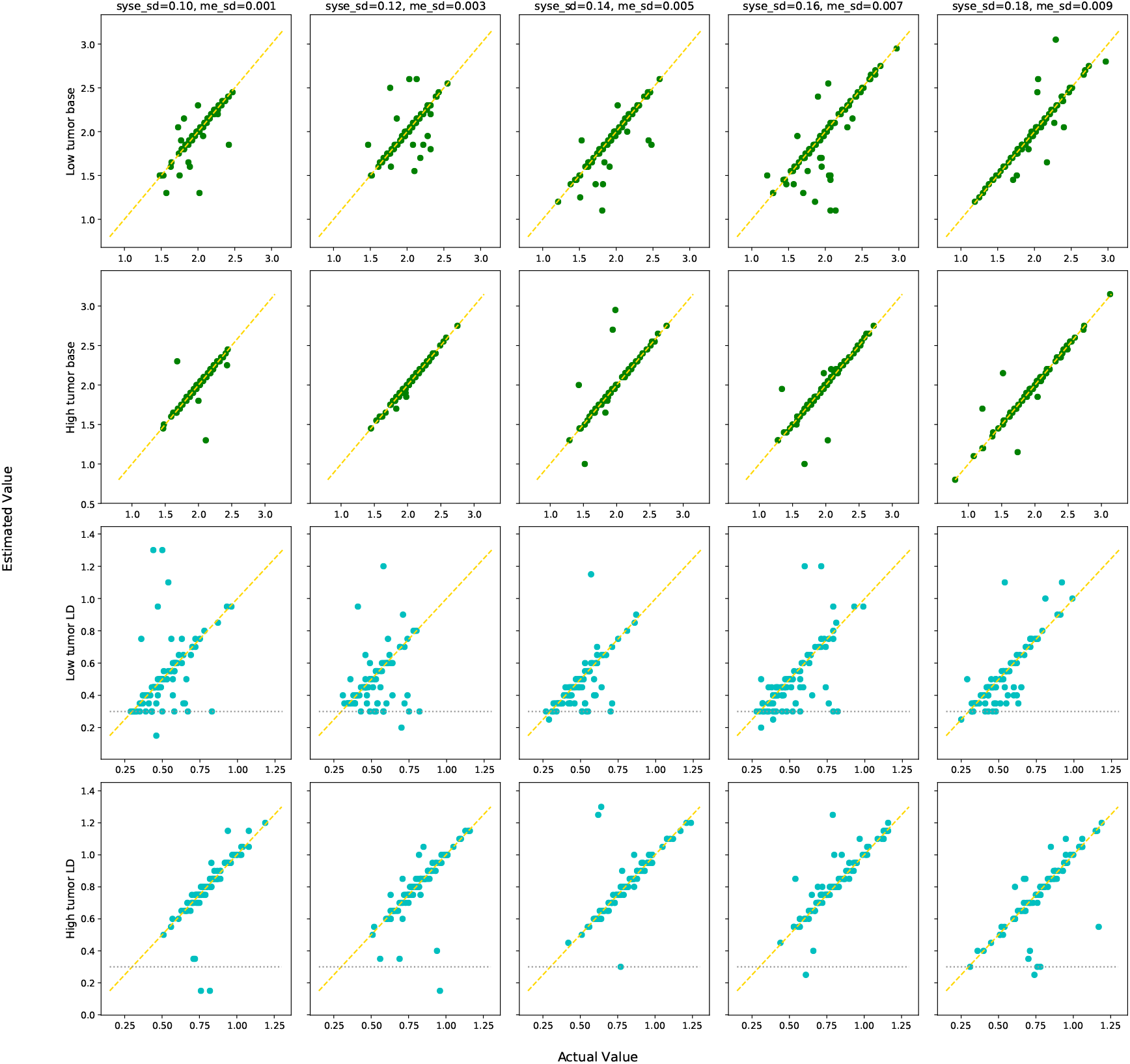
The comparisons of estimated values and actual values on simulated data with different settings. The first row shows results of baseline estimation in low tumor composition; the second row shows results of baseline estimation in high tumor composition, the third row shows results of level distance estimation in low tumor composition, and the fourth row shows results of level distance estiation in high tumor composition. The noise level increases gradually in each column from left to right. The dotted lines indicate a threshold that can filter out problematic results.

Overall *Mecan4CNA* performed with high accuracy and consistency in all scenarios. Specifically, for the estimation of baseline, *Mecan4CNA* achieved 97% average accuracy in the combination of all high tumor composition datasets; 89% average accuracy in the combination of all low tumor composition datasets; and 93% overall accuracy of all datasets. For the estimation of level distance, *Mecan4CNA* achieved 96% average accuracy in the combination of all high tumor composition datasets; 86% average accuracy in the combination of all low tumor composition datasets; and 91% overall accuracy of all datasets.

### Applications of *Mecan4CNA*

#### Baseline deviation of TCGA data

The Cancer Genome Atlas (TCGA) provides a large collection of copy number data of from 33 cancer projects. The copy number data was generated using Affymatrix SNP6 microarrays and a uniform processing pipeline. We used *Mecan4CNA* to estimate the baseline value of all TCGA copy number data, summarized by project. As shown in Figure 3, baseline deviation is widespread with varying contributions related to individual projects. In general, the deviation level is strongly correlated with the difficulty of acquiring pure biopsies. In projects such as LAML (Acute Myeloid Leukemia) and CESC (Cervical Squamous Cell Carcinoma and Endocervical Adenocarcinoma), where biopsies are usually pure, the original baseline levels are often accurate and the deviation is often low. In projects such a LUSC (Lung Squamous Cell Carcinoma) and OV (Ovarian Serous Cystadenocarcinoma), where biopsy are difficult to acquire and frequently contain heterogeneous cellular populations, the original baseline levels often have high deviation.

**Figure 3:**
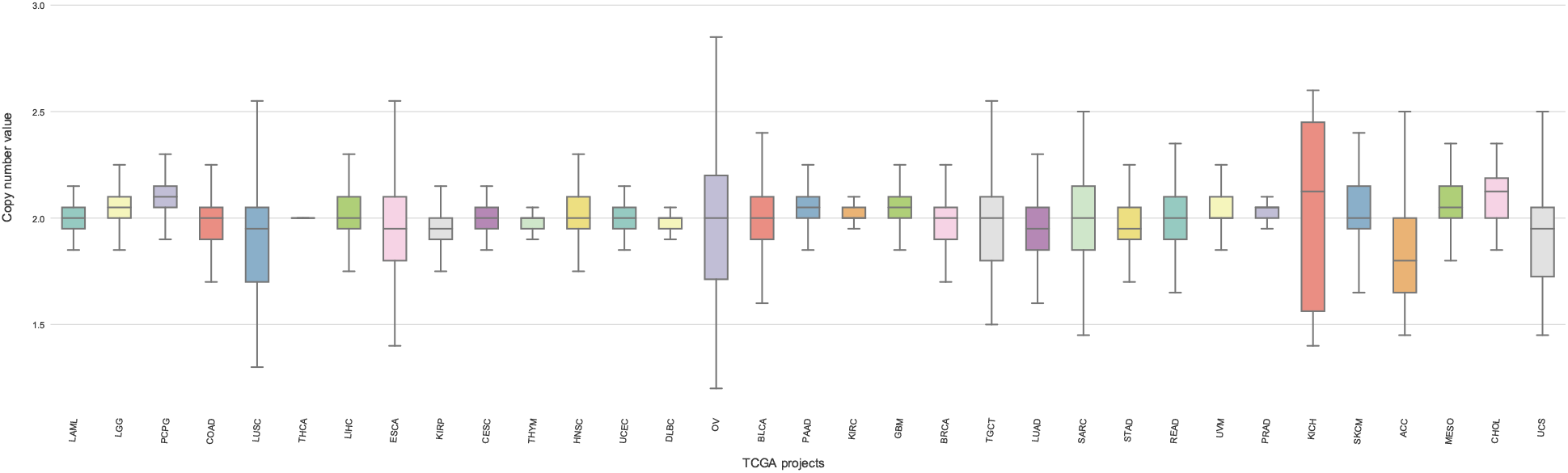
The distribution of baseline values estimated with *Mecan4CNA* in different TCGA projects. Outliers are not shown.

Although great efforts went into limiting errors and noise in TCGA copy number data, they originated from projects addressing diverse diseases and where data generation were different in time, space and experimental conditions. The variation in the results does not directly reflect on the accuracy of the results since the determination of an underlying “Gold Standard” is beyond the scope of our methodology. The generally good quality of these datasets allowed us to show that baseline variation is a common and recurring problem among copy number data.

#### Normalized data as GISTIC input

In order to demonstrate the effect of *Mecan4CNA* in copy number data analysis, we applied GISTIC with both the original and the normalized copy number data from the TCGA project. GISTIC is a widely used multi-sample tool to detect and score focal regions in copy number datasets. We chose data from 3 TCGA projects with different baseline deviation levels to look for focal regions and potential driver genes using GISTIC. GISTIC was run in *gene-gistic* mode with default parameters. Data was normalized using the following formula:

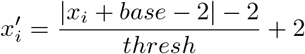

Where *base* is the estimated baseline value and *thresh* is *level distance*.

Table 1 illustrates the comparisons of called regions from GISTIC using the original and normalized data. Overall, when using the normalized data, the number of detected regions is significantly reduced (Figure 4). At the same time, when mapping focal regions to Cancer Census Genes [20], the number the detected drivers actually increases as shown in Figure 6. Some driver genes are only captured using the normalized data, such as PDGFRA in GBM, XPC in SKCM and CASP3 in OV. For focal regions detected by both settings, the size of the region is usually reduced by the normalized data (Figure 5). A few new focal regions containing driver genes, which were too weak in signal when using the original data, became significant and were detected using the normalized data. For example, in the SKCM project region *chr6:9090525-9120556*^2^ was detected by the original data with high significance but harbors no protein coding genes. When using the normalized data, the signal strength of this region was reduced dramatically and it was not called as a focal region. Region *chr12:68834906-68893964* was called only when using the normalized data and harbors two known oncogenes (CMP and MDM2). Region *chr1:204403604-204437602* was narrowed and shifted to *chr1:204477206-204586780* by using the normalized data, this subtle change removed a passenger gene PPP1R15B and included a oncogene MDM4. The Molecular Signatures Database (MSigDB) [11] is a collection of annotated gene sets for use with GSEA software. The “C6: oncogenic signatures” represent signatures of cellular pathways which are often dis-regulated in cancer. We mapped the genes called by GISTIC using the original and the normalized data to each pathway in C6. Table 2 shows pathways that are covered by a high number of genes in both data settings. Pathways in bold are with known associations to melanoma. Typically, Both KRAS and P53 are prevalent and key driver genes in melanoma. Although these pathways stood out in both data settings, most of these pathways were covered with more genes when using the normalized data. Table 3 shows pathways with significant coverage difference between the original and the normalized data. The coverage of several pathways (KRAS, PTEN, MYC), which have strong associations to tumorigenesis in melanoma, were significantly increased when using the normalized data. In summary, when using the normalized data instead of the original data, the analysis result of GISTIC showed a prominent improvement in both sensitivity and specificity.

**Table 1:** Summary of detected regions of GISTIC using original and normalized data. Numbers in column represents the count of gain/loss.

|  | Data | No. regions | No. genes | No. Census genes | Gene density | Driver density |
| --- | --- | --- | --- | --- | --- | --- |
| GBM | Original | 29/46 | 143/1722 | 19/45 | 4.93/37.43 | 0.13/0.03 |
|  | Normalized | 26/40 | 110/1953 | 19/50 | 4.23/48.83 | 0.17/0.03 |
| SKCM | Original | 33/37 | 1494/1721 | 58/43 | 45.27/46.51 | 0.04/0.02 |
|  | Normalized | 26/21 | 1860/2099 | 47/66 | 71.54/99.95 | 0.03/0.03 |
| OV | Original | 38/51 | 423/2515 | 13/64 | 11.13/49.31 | 0.03/0.03 |
|  | Normalized | 35/36 | 382/2240 | 13/54 | 10.91/62.22 | 0.03/0.02 |

**Figure 4:**
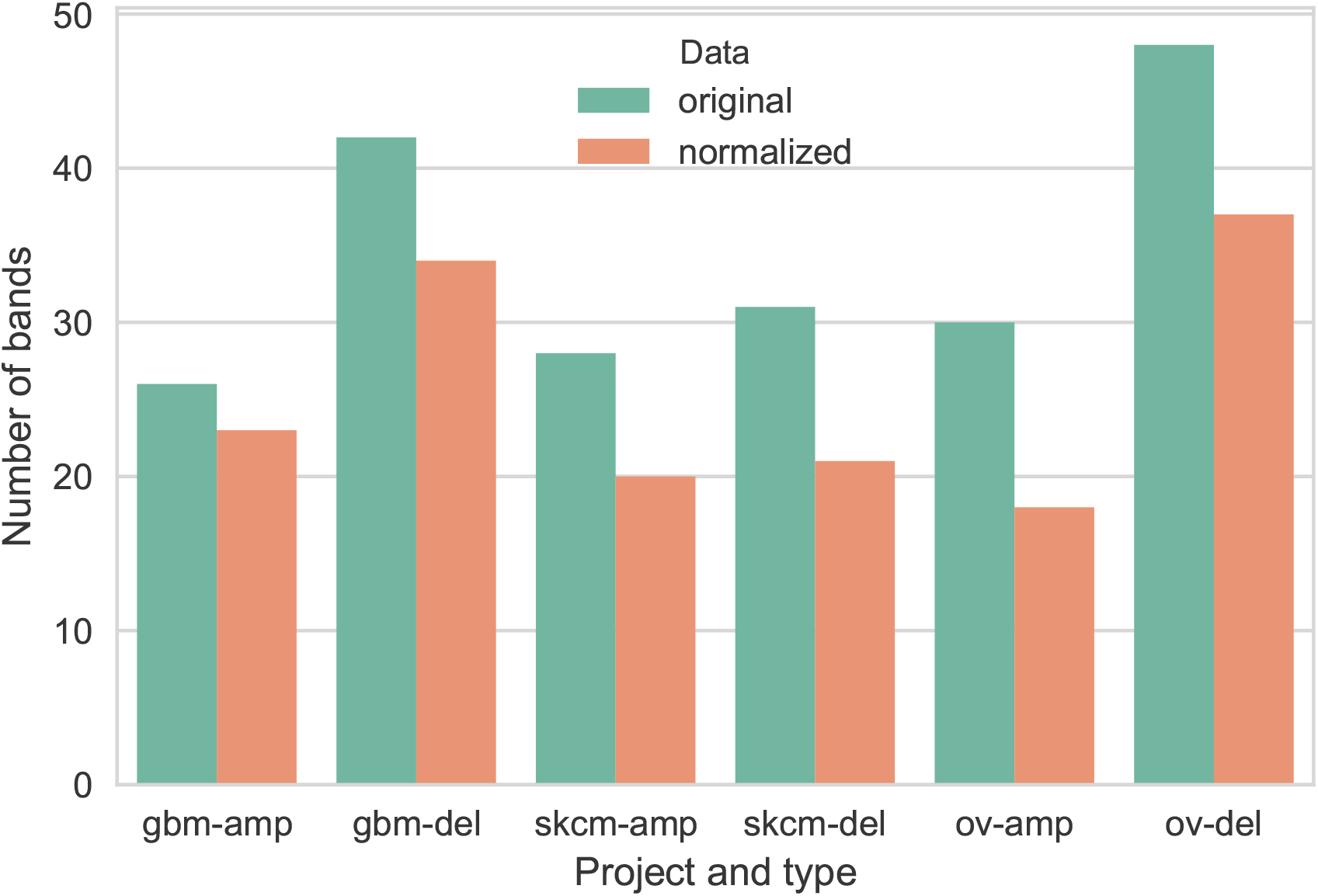
The number of significant bands detected by GISTIC using different datasets.

**Figure 5:**
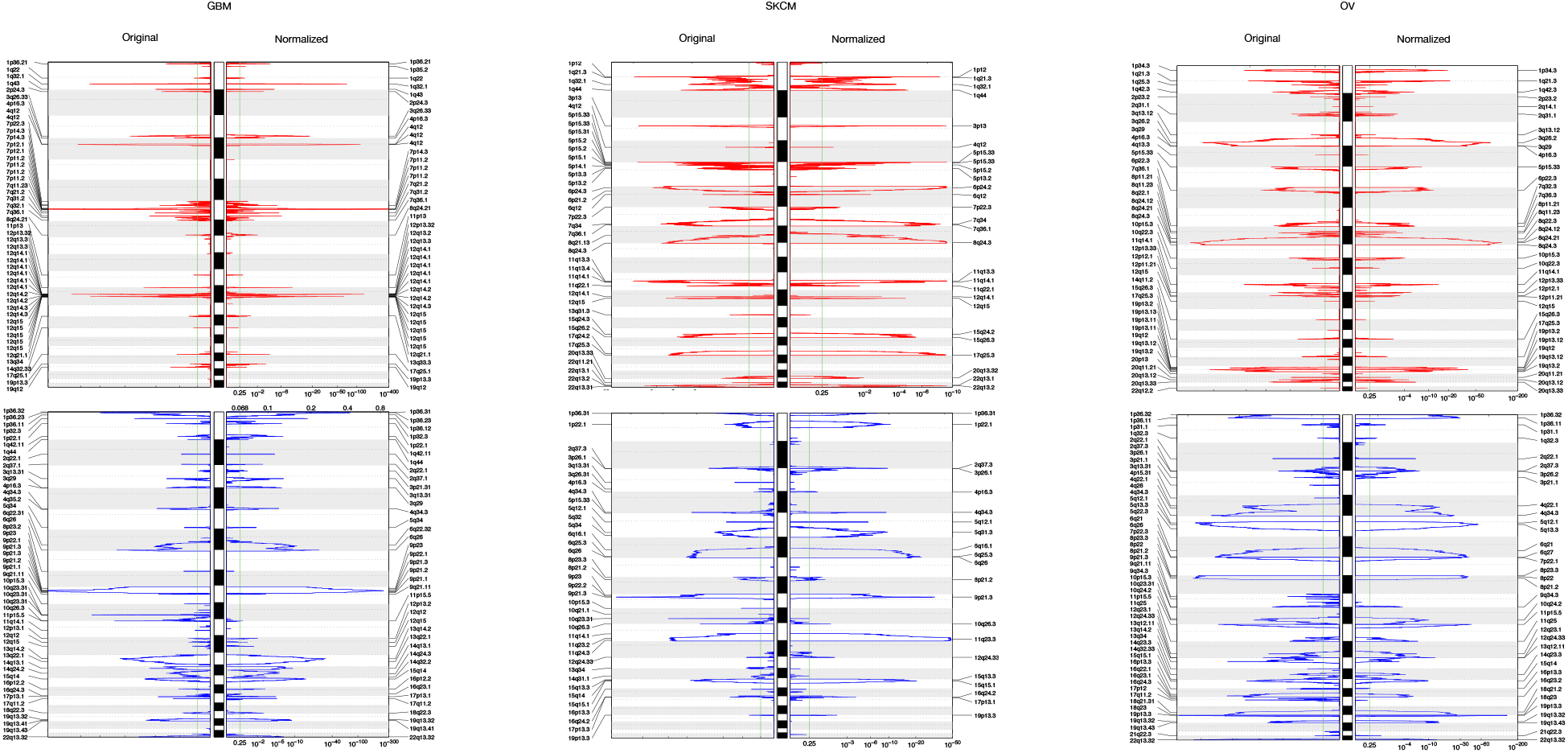
The comparisons of GISTIC calling results using the original and normalized copy number data from 3 TCGA projects. The green boxes highlight regions with reduced noise signal using normalized data. The orange boxes highlight regions with improved significance using normalized data. Overall, normalized data shows improvement in both sensitivity and specificity.

**Figure 6:**
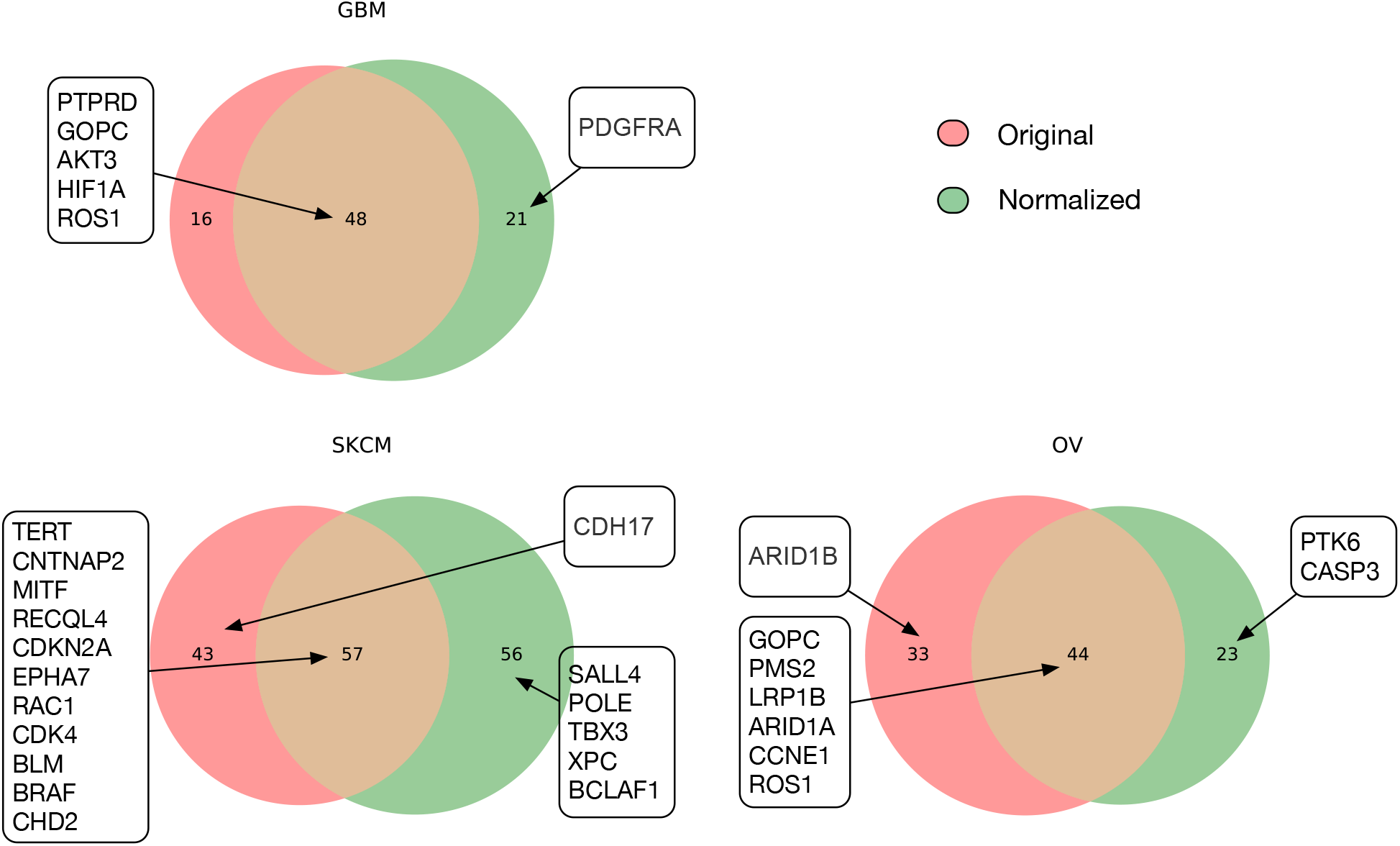
Detected cancer census genes using the original and normalized data in GBM, SKCM and OV datasets of TCGA. Numbers in circles represents the total of detected census genes. Gene symbols in boxes show the known drivers for the disease.

**Table 2:**
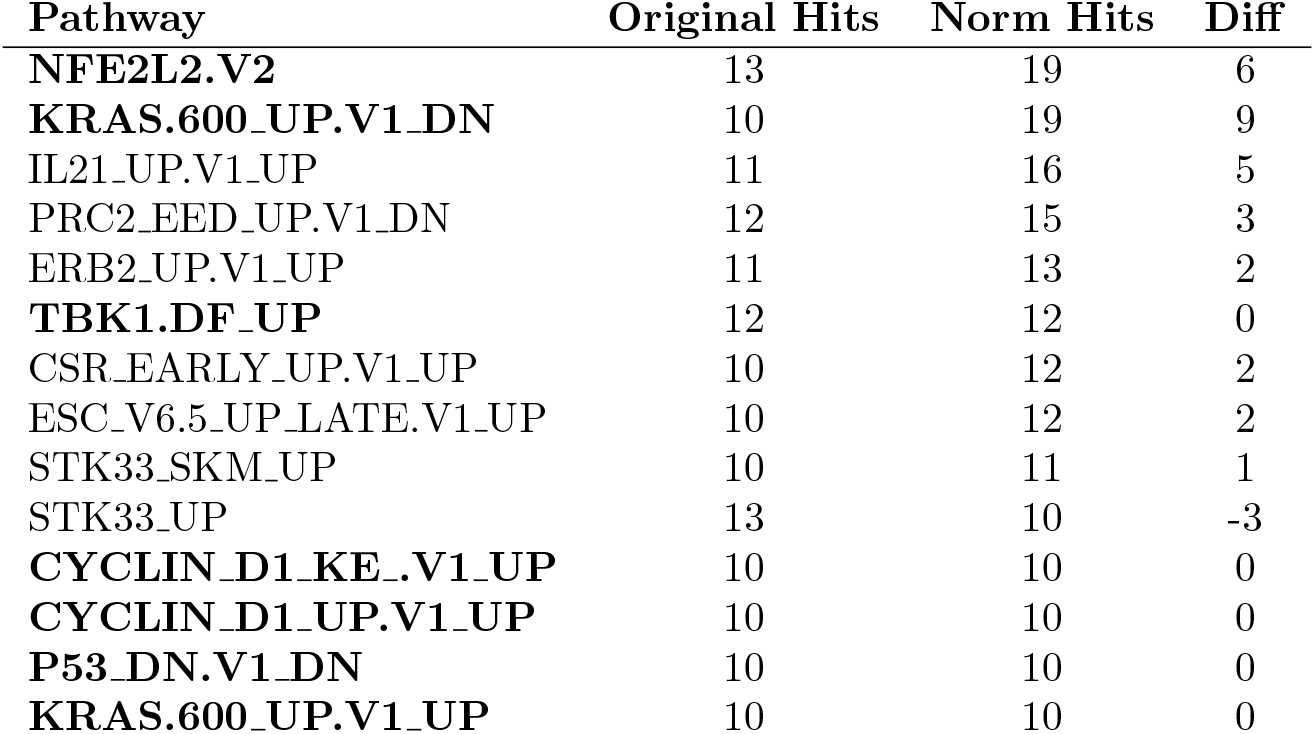
MSigDB pathways with high coverage in both the original and normalized data of SKCM. Pathways in bold have known associations with melanoma.

**Table 3:** MSigDB pathways with significant changes of coverage between the original and normalized data of SKCM. Pathways in bold have known associations with melanoma.

| Pathway | Original Hits | Norm Hits | Diff |
| --- | --- | --- | --- |
| PDGF_ERK.DN.V1.DN | 3 | 15 | 12 |
| NRL.DN.V1.UP | 1 | 11 | 10 |
| **KRAS.600.UP.V1.DN** | 10 | 19 | 9 |
| IL2.UP.V1.UP | 8 | 17 | 9 |
| VEGF_A.UP.V1.UP | 7 | 16 | 9 |
| STK33.DN | 3 | 12 | 9 |
| CRX_NRL.DN.V1.UP | 1 | 10 | 9 |
| MTOR.UP.V1.UP | 5 | 13 | 8 |
| PIGF.UP.V1.DN | 4 | 12 | 8 |
| STK33.NOMO.DN | 2 | 10 | 8 |
| TGFB.UP.V1.UP | 7 | 15 | 8 |
| RPS14.DN.V1.DN | 7 | 15 | 8 |
| **KRAS.300.UP.V1.DN** | 5 | 12 | 7 |
| **KRAS.AMP.LUNG.UP.V1.DN** | 2 | 9 | 7 |
| PDGF.UP.V1.UP | 6 | 13 | 7 |
| RAPA_EARLY.UP.V1.UP | 8 | 15 | 7 |
| IL15.UP.V1.UP | 4 | 11 | 7 |
| MTOR.UP.V1.DN | 3 | 10 | 7 |
| **PTEN.DN.V1.UP** | 9 | 16 | 7 |
| MEK.UP.V1.UP | 7 | 14 | 7 |
| **MYC.UP.V1.UP** | 5 | 12 | 7 |
| CAHOY_ASTROCYTIC | 1 | 8 | 7 |
| MEL18.DN.V1.DN | 9 | 4 | -5 |
| PRC2_EZH2.UP.V1.DN | 10 | 3 | -7 |

## Conclusion

In this study, we presented a novel method dubbed Minimum Error Normalization of Copy Numbers (Mecan4CNA). When tested on simulation data and cell line data, it showed promising accuracy in estimating the baseline and the level distance of a copy number profile. When applied on the copy number data from TCGA, it illustrates that baseline deviation widely exists among copy number profiles and is primarily attributed to the purity of the sample. Finally, the comparison of results from GISTIC, when using the original and the normalized data, showed that the normalized data can provide a significant improvement in both sensitivity and specificity. *Mecan4CNA* does not rely on any additional information nor prior knowledge, and is also efficient in speed and resource. It is designed to analyze and normalize multiple samples, but can also be used for single sample interpretation.

## Software availibility

1. The pip version: https://pypi.org/project/mecan4cna/
2. Latest source code: https://github.com/baudisgroup/mecan4cna
3. Software license: MIT

## Footnotes

1 Alternatively to “Copy Number Aberrations” (CNA) the term “Somatic Copy Number Variations” (sCNV) is being used in the literature occasionally. We prefer the CNA term due to precedence and epistemic relation to the oncogenetic process.

2 base positions according to reference genome GRCh38

